# Chromosome length and gene density contribute to micronuclear membrane stability

**DOI:** 10.1101/2021.05.12.443914

**Authors:** Anna E Mammel, Heather Z Huang, Amanda L Gunn, Emma Choo, Emily M Hatch

## Abstract

Micronuclei are derived from missegregated chromosomes and frequently lose membrane integrity, leading to DNA damage, innate immune activation, and metastatic signaling. Here we demonstrate that two characteristics of the trapped chromosome, length and gene density, are key contributors to micronuclei membrane stability in human cells. Chromosome length is proportional to micronuclei size, and gene density has an additive effect with micronucleus size on membrane stability. We demonstrate that these results are not due to chromosome-specific differences in spindle position or initial nuclear pore complex recruitment during post-mitotic nuclear envelope assembly. We find that chromosome length and micronuclei size strongly correlate with lamin B1 and nuclear pore density in intact micronuclei. Unexpectedly, lamin B1 levels do not predict nuclear lamina organization and membrane stability. Instead, small gene-dense micronuclei have decreased nuclear lamina gaps compared to large micronuclei, despite very low levels of lamin B1. Our data strongly suggest that nuclear envelope composition defects previously correlated with membrane rupture only partly explain membrane stability in micronuclei. We propose that an unknown factor linked to gene density has a separate function that inhibits the appearance of nuclear lamina gaps and delays membrane rupture until late in the cell cycle.

## Introduction

Micronuclei (MN) in metazoans form around chromosomes or chromosome fragments that missegregate during mitosis and recruit their own nuclear envelope (NE). MN are biomarkers of chromosome instability in cancer, frequently arise during early embryogenesis in humans, and occur at a low frequency in healthy tissue (Guo et al., 2018). Similar to nuclei, MN are enclosed by a nuclear membrane and typically have a nuclear lamina and nuclear pores, although often at a reduced density (Hatch et al., 2013; Liu et al., 2018). MN can support major nuclear functions, including transcription, DNA replication, and DNA damage repair, although these are often attenuated or delayed (Crasta et al., 2012; Hatch et al., 2013; Liu et al., 2018; Terradas et al., 2012, 2009). However, persistent rupture of the MN membrane, which causes loss of MN compartmentalization for the duration of interphase, occurs at a high frequency in cultured cells, cancer tissue, and early embryogenesis (Daughtry et al., 2019; Hatch et al., 2013; Liu et al., 2018; Vázquez-Diez et al., 2016; Willan et al., 2019). MN rupture arrests micronuclear functions and leads to aneuploidy, DNA damage, and activation of innate immune and cell invasion pathways (Bakhoum et al., 2018; Harding et al., 2017; Hatch et al., 2013; Ly et al., 2016; Mackenzie et al., 2017; Mohr et al., 2020; Soto et al., 2018; Zhang et al., 2015). DNA damage caused by MN rupture is thought to be a major driver of chromothripsis and kataegis, two “all-at-once” processes that cause chromosomal rearrangements and hyper-mutations, respectively (Nik-Zainal et al., 2012; Stephens et al., 2011). A current model is that MN rupture causes fragmentation of the encapsulated chromatin, which remains together through mitosis and is re-ligated by error-prone DNA damage repair pathways upon incorporation into a nucleus or micronucleus in the next cell cycle (Crasta et al., 2012; Hatch et al., 2013; Ly et al., 2019, 2016; Umbreit et al., 2020; Zhang et al., 2015). Despite the high frequency of MN rupture and its potential to drastically change gene expression, the molecular mechanisms of membrane rupture in MN and its consequences are unclear.

MN frequently have large gaps in the nuclear lamina meshwork that leave areas of weak membrane that are the site of membrane rupture (Hatch et al., 2013). Similar gaps enable and are the site of membrane rupture in the nucleus (Maciejowski and Hatch, 2020). Lamin B1 depletion is sufficient to cause nuclear lamina gap formation in nuclei (Hatch and Hetzer, 2016; Lenz-Böhme et al., 1997; Shimi et al., 2008; Vergnes et al., 2004) and impaired lamin B1 recruitment is thought to underlie defects in nuclear lamina organization and membrane stability in MN as well (Hatch et al., 2013; Kneissig et al., 2019; Liu et al., 2018; Okamoto et al., 2012; Xia et al., 2019). Overexpression of lamin B1, or its related protein lamin B2, is sufficient to inhibit nuclear membrane rupture in both MN and nuclei (Bakhoum et al., 2018; Hatch et al., 2013; Hatch and Hetzer, 2016; Maciejowski et al., 2015; Vargas et al., 2012). However, nuclear lamina gaps can form many hours before the MN loses integrity (Hatch et al., 2013), suggesting that other mechanisms trigger membrane disruption. Actomyosin compression likely accelerates rupture of very large MN (Liu et al., 2018), but the trigger in most cases is unknown (Hatch and Hetzer, 2016).

A current model for reduced NE protein recruitment to MN is that the microtubule-dense midspindle prevents targeting of critical components, including lamin B1 and nuclear pore complexes (NPCs), to lagging chromosomes during NE assembly by inhibiting protein dephosphorylation or physically impairing ER access (Afonso et al., 2014; Castro et al., 2017; Karg et al., 2015; Liu et al., 2018). In these models, chromosomes missegregating outside the spindle or at the spindle poles generate larger MN that recruit near normal amounts of lamin B1 and NPCs and have a lower frequency of membrane rupture (Liu et al., 2018). Other data suggest high membrane curvature and small nuclear size are sufficient to impair lamin B1 meshwork assembly in both the nucleus and MN (Kneissig et al., 2019; Xia et al., 2019, 2018).

The composition and stability of the nuclear lamina could vary widely between single chromosome MN depending on the identity of the entrapped chromosome. Heterochromatin is a key regulator of nuclear lamina organization, nuclear mechanical stability, and nuclear membrane integrity, and its density varies widely between chromosomes in the human karyotype (Furusawa et al., 2015; Stephens et al., 2018). Chromosome identity could also contribute to nuclear lamina composition in MN, since NE proteins bind to heterochromatin histone modifications directly, interact with the nuclear lamina, and regulate NE assembly and chromosome location in interphase (Pyrpasopoulou et al., 1996; Solovei et al., 2013; Zullo et al., 2012). Differences in chromosome length and centromere size between individual chromosomes could also indirectly affect MN nuclear lamina recruitment by biasing chromosome position to outside or within the midspindle during missegregation (Booth et al., 2016; McIntosh and Landis, 1971; Mosgöller et al., 1991). Thus, chromosome identity could contribute to MN stability through multiple mechanisms.

In this paper we demonstrate that chromosome identity is a major determinant of MN rupture timing. Analysis of single chromosome MN finds that chromosome length, which correlates linearly with MN size, and gene density have an additive effect on membrane integrity and delay membrane rupture. Chromosome-based MN stability differences are not due to a bias in missegregation positioning. Chromosomes correlated with high and low MN stability have similar midspindle missegregation localization and similar NPC recruitment defects during post-mitotic NE assembly, suggesting that differences occur at a later time point. Instead, we find a strong correlation between lamin B1 levels, NPC density, and MN size in interphase. Surprisingly, small gene-dense MN have very low levels of lamin B1 but are less likely to have nuclear lamina gaps compared to gene-poor MN of similar size, suggesting that gene density is a strong predictor of nuclear lamina organization. Our data confirm a connection between MN size and nuclear lamina composition, but indicate that an intrinsic factor linked to high gene density is sufficient to inhibit nuclear lamina disorganization even in the absence of lamin B1. Together, these results signify that analyzing MN chromosome content will be critical to understanding the cellular consequences of micronucleation in different disease contexts.

## Results

To analyze chromosome-specific differences in MN stability, we first established a robust system to identify single micronucleated chromosomes by fluorescent *in situ* hybridization (FISH), using commercially available *<u>H</u>omo <u>sa</u>piens* (HSA) chromosome specific probes, combined with immunofluorescence (IF) against a centromere protein. Single chromosome MN were generated in hTERT-RPE-1 cells, a near-diploid chromosomally stable cell line, by first synchronizing these cells in G1 with a cdk4/6 inhibitor (PD-0332991) then releasing cells into reversine (mitotic kinases monopolar spindle 1 inhibitor, Mps1i), which inhibits the spindle assembly checkpoint (Santaguida et al., 2010) (Fig. 1A). MN rupture frequency was assessed by histone H3K27 acetylation (H3K27ac) IF. We previously demonstrated that histone acetylation marks are lost upon MN rupture (Hatch et al., 2013). To validate H3K27ac as a sensitive and accurate marker of intact MN in our system, we quantified the co-localization of H3K27ac and 3xGFP-NLS (nuclear localization signal) in MN and identified a strong correlation between the two marks (Fig. S1A, B). Only a small proportion of MN were H3K27ac positive, 3xGFP-NLS negative, suggesting that the majority of MN assembled NPCs and had nuclear transport. In addition, we observed a similar decrease in the number of H3K27ac positive MN during interphase (Fig. S1C, D), as had been observed previously using GFP-NLS (Hatch et al., 2013; Liu et al., 2018; Zhang et al., 2015), suggesting that H3K27Ac accurately labeled intact MN. To validate H3K27ac as an integrity marker for MN containing small gene-poor chromosomes, we assessed H3K27ac labeling of single chromosome HSA 18 MN. HSA 18 MN frequently had reduced H3K27ac labeling compared to the nucleus, but the signal was sufficiently high to distinguish intact from ruptured MN (Fig. 1B). This was true for all other chromosomes examined in this study (Fig. S1E, F).

**Figure 1.**
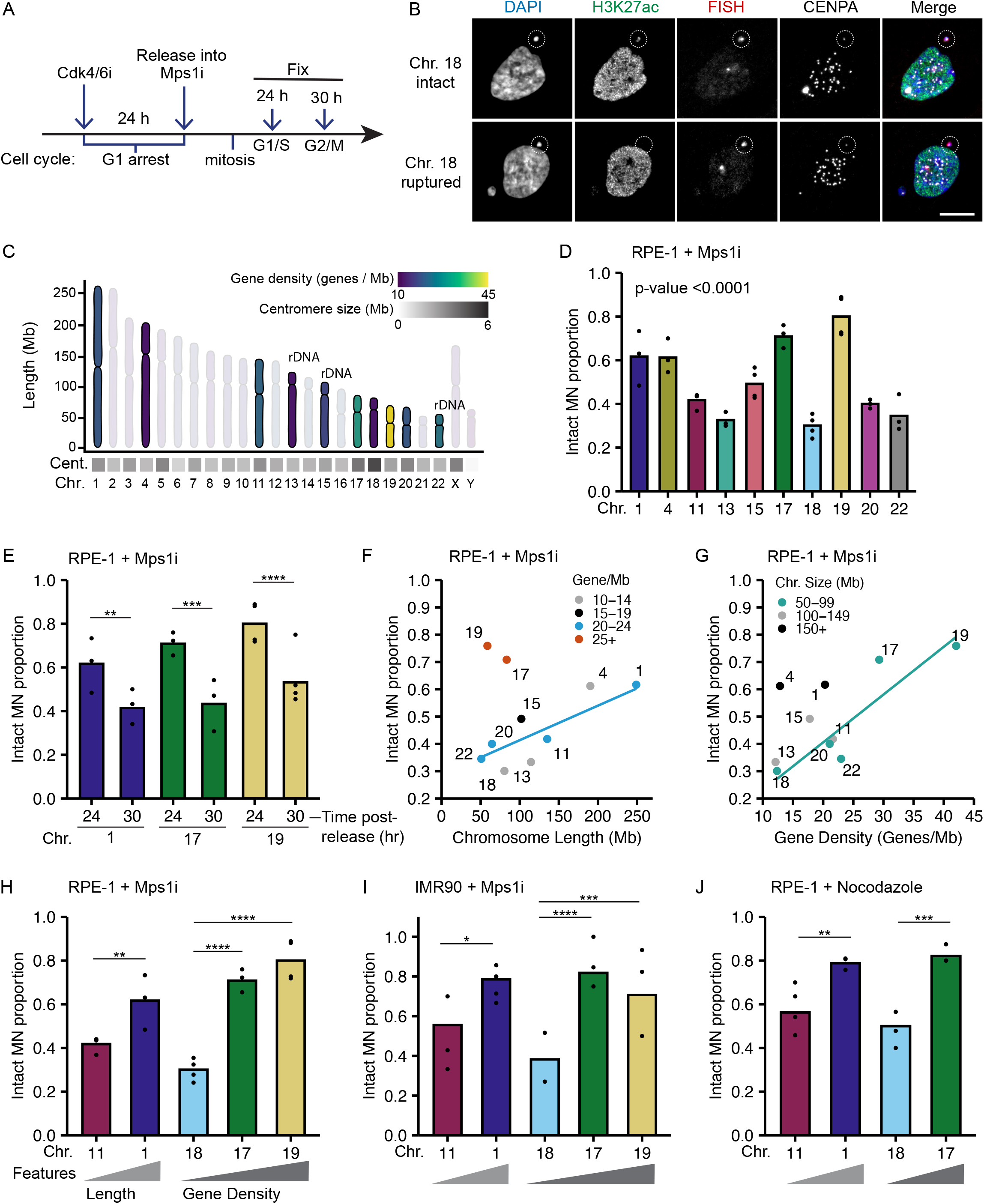
Micronucleated chromosome length and gene density correlate with membrane rupture. (**A**) Time course of IF/FISH experiments. RPE-1 cells arrested in G1 by incubation in PD-0332991 (Cdk4/6i) for 24 hours, then released into 0.5 μM reversine (Mps1i) to induce MN formation. Cells fixed 24 or 30 hours post-Cdk4/6i release, at G1/S and S/G2, respectively (see Fig. S1G, H). (**B**) Maximum intensity projection (MIP) images of intact (H3K27ac+) and ruptured (H3K27ac-) MN containing a single HSA 18 (dotted circle). Scale bar 10 μm. (**C**) Characteristics of selected chromosomes (bold) represent a wide range of chromosome length, centromere size and positioning, rDNA repeats, and gene density. (**D**) Intact proportion 24 hours post-Cdk4/6i release for single chromosome containing MN. Chi-square; p-value < 0.0001; N = 3 – 4; n = 67 – 133. (**E**) Comparison of MN rupture frequency 30 h to 24 h post-release (from panel D), N = 3 – 4; n = 60 – 111. (**F – H**) Replotting of data from panel D. (F) MN stability (proportion intact) positively correlates with chromosome length when grouped by gene density, (G) and with gene density, when grouped by chromosome length. Only groups with > 3 chromosomes were analyzed (line). (**H**) MN stability correlates with chromosome length and gene density for representative chromosomes: HSA -1 compared to -11 for length, and HSA 18 compared to HSA -17 and -19 for gene density. (**I**) MN stability in single chromosome MN in IMR90 cells treated as in panel A. Similar correlations observed between MN stability and chromosome length and gene density as in panel H. N = 3 – 4; n = 33 – 68. (**J**) MN stability in single chromosome MN in RPE-1 cells after nocodazole release. Similar correlations observed between MN stability and chromosome length and gene density as in panel H. N = 3 – 4; n = 56 – 85. For all bar graphs in paper, individual experiments are represented as points and pooled replicate proportions are represented as bars. For all sample sizes, N = number of experimental replicates, n = number of objects per replicate. Chi-square tests are performed for all experiments prior to pairwise comparisons by Barnard’s exact test, except where indicated. * p < 0.05, ** p < 0.01, *** p < 0.001, **** p < 0.0001. Cent. = centromere, Chr. = chromosome, Mb = megabase.

To determine whether chromosome identity correlated with MN stability, and which chromosome features regulated membrane integrity, we examined a panel of 10 chromosomes spanning the distribution of chromosome length (5 fold, [*NIH – Human genome assembly GRCh38*.*p13*]), gene density (3.5 fold, (Worrall et al., 2018)), centromere size (4.2 fold, (Miga et al., 2014)), rDNA presence, and centromere position in the human karyotype (Fig. 1C, Fig. S1F). Rupture frequency was compared between different single-chromosome MN 24 hours post-release in Mps1i (Fig. 1A) when approximately 50% of MN were ruptured (Fig. S1D) and cells were in G1/ S (Fig. S1G, H). We found consistent chromosome-specific differences in MN stability across multiple experimental replicates (Fig. 1D), with several chromosomes having a high likelihood of maintaining MN stability throughout G1. Analysis of MN rupture frequency of highly intact chromosome MN at a later time point in S/G2 (Fig. S1 G, H), found that chromosome identity delays but does not prevent rupture (Fig. 1E).

Examination of the traits most closely correlated with MN stability identified chromosome length and gene density as directly proportional to MN integrity. HSA -1, -11, -20, and -22 have a similar gene density (20 – 23 genes/Mb) but vary 5-fold in length, and the proportion of intact MN (MN stability) consistently increases with increasing length (Fig. 1F, H). Similar results are observed for HSA -4, -13, and -18, which have a lower gene density (12 – 13 genes/Mb) and vary 2-fold in length (Fig. 1F). Surprisingly, comparison of HSA -17, -18, -19, -20, and -22 which have similar length (50 - 100 Mb) but a 3.4-fold variation in gene density, showed a consistent increase in MN stability with increasing gene density (Fig. 1G). Similar trends were not observed for centromere size (Fig. S2A), and MN stability was not altered by the presence of rDNA or acrocentric centromeres when compared to chromosomes of similar length and gene density (Fig. S2B, C). To determine whether these correlations between gene density or chromosome length and stability were conserved between MN formation mechanisms and cell lines, we assessed the rupture frequency of HSA -1 vs. -11 (chromosome length) and HSA -18 vs -17 and/or -19 (chromosome gene density) using an alternative Mps1i (BAY-1217389), after nocodazole release, which increases prometaphase duration, and in IMR90 fibroblasts treated with Mps1i (reversine). In each case, we found that MN containing larger or more gene dense chromosomes were more stable (Fig. 1H-J, Fig. S2D), suggesting that these correlations are independent of the mitotic defect or gene expression specific to RPE-1 cell identity.

Chromosome volume in the nucleus is proportional to the length of the chromosome (Eils et al., 1996; Kemeny et al., 2018) and we identified a similar relationship between chromosome length and MN area (Fig. 2A). The relationship was best fit by linear regression, rather than an exponential equation, because MN in RPE-1 cells were closer to oblate spheroids than spheres across all MN (Fig. 2B, C), consistent with a strong correlation between MN area and volume (Fig. S3A). To test the hypothesis that MN size determines rupture frequency, we induced multi-chromosome MN by treating with a higher dose of Mps1i (Fig. S3B). Both the median MN area and the proportion of intact MN increased with centromere number (Fig. 2D, E), consistent with increased size improving stability. However, we found that even very large multi-chromosomal MN were not protected from membrane rupture later in the cell cycle (Fig. 2E, Fig. S3C).

**Figure 2.**
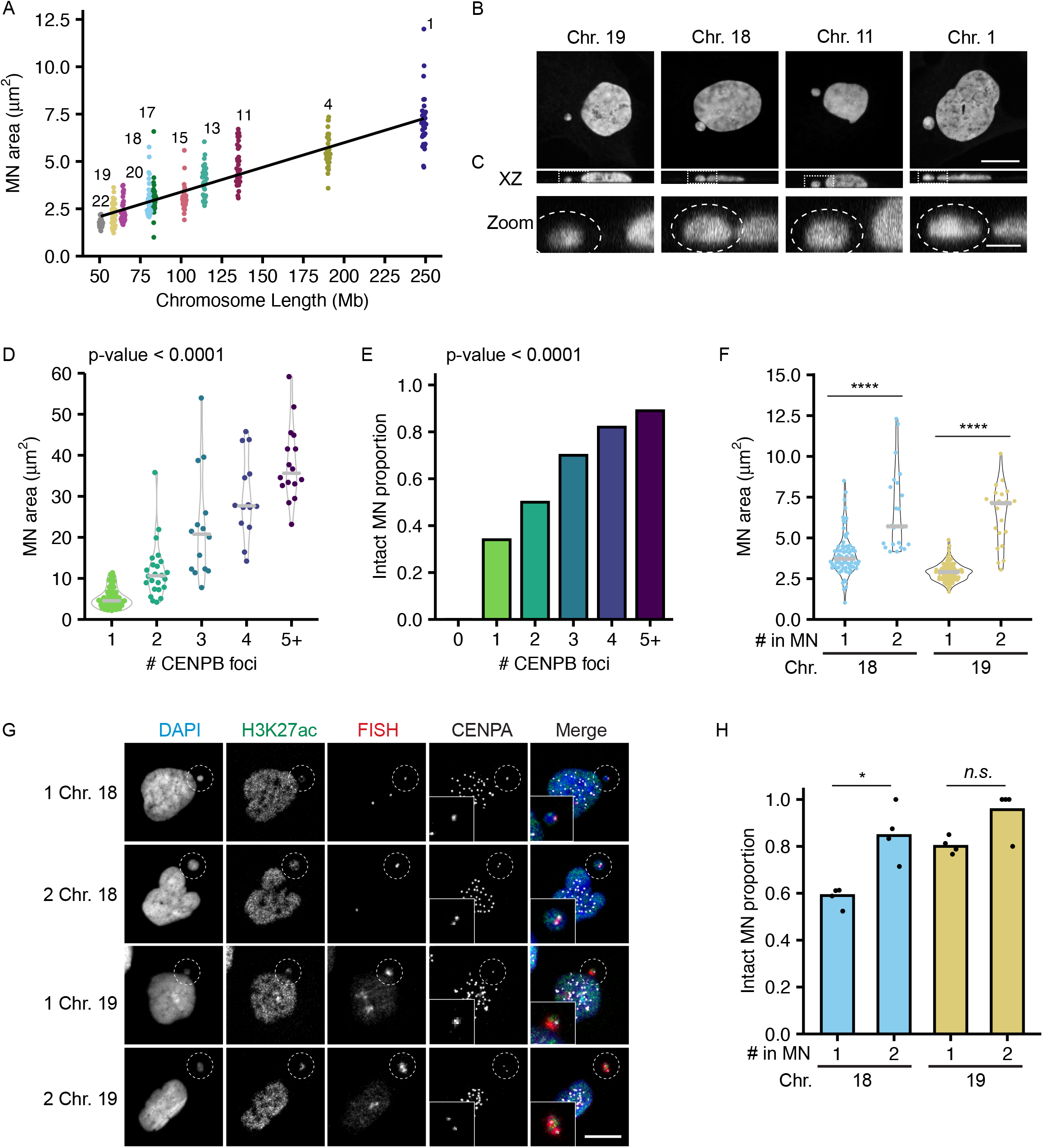
MN size correlates with MN stability and has an additive effect to gene density. (**A**) Maximum projected area of intact MN 24 h post-Cdk4/6i release is linearly correlated with chromosome length in RPE-1 cells (see Fig. S1). Spearman correlation of mean MN area; R^2^ = 0.76, **** p < 0.0001, n = (25, 30, 33, 30, 38, 30, 28, 30, 31, 30) MN per chromosome. (**B**) Representative images of DAPI stained intact single chromosome MN containing indicated chromosomes (FISH/CENPA labeling not shown). Orthogonal sections (XZ planes) shown for each. Scale bar = 10 μm. (**C**) Zoomed in image of the XZ orthogonal section (white box) containing the MN, with DAPI outlined. Z-step size = 0.15 μm; 21 steps; scale bar = 2 μm. (**D - E**) Quantification of intact MN area (D) and MN stability (E) in RPE-1 cells 24 hours post-Cdk4/6i release into 1μM MPS1i. MN containing single and multiple chromosomes, determined by CENPB foci number, were analyzed. All MN lacking a centromere were ruptured, and therefore not included in area analysis. (D) Kruskal-Wallis one-way test; p-value < 0.0001; N = 3, n = (61, 24, 14, 14, 17). (E) Chi-square; p-value < 0.0001; N = 3, n = (10, 177, 48, 20, 17, 19). (**F**) MN containing two copies of HSA -18 or -19 are significantly larger than MN containing a single copy. Welch two sample t-test; **** p-value < 0.0001; N = 4, n = (72, 22, 92, 22). (**G**) Images of intact (H3K27ac+) MN containing one or two copies of HSA -18 and -19. Chromosome number determined by CENPA foci number. Scale bar = 10 μm. (**H**) MN containing two copies of HSA 18 had significantly increased stability compared to MN containing a single copy. MN containing two copies of HSA 19 were more likely to be intact than MN containing a single copy, but the difference was not statistically significant. Barnard’s exact test; * p < 0.05, *n*.*s*. p > 0.05. N = 4, n = (122, 26, 115, 23).

To determine the relationship between MN size and gene density, we assessed MN rupture frequency in MN containing either one or two copies of HSA 18 (gene-poor) or HSA 19 (gene-dense) (Fig. 2F – H). For both chromosomes, doubling the number of alleles increased the median area (Fig. 2F, G). Increasing MN size rescued the membrane instability of the gene-poor HSA 18 MN and further increased the stability of the gene-dense HSA 19 MN (Fig. 2H). These data indicate that MN size and gene density are additive and suggest that they regulate MN stability through independent mechanisms.

Larger chromosomes tend to segregate on the exterior of the metaphase plate during mitosis (Booth et al., 2016; McIntosh and Landis, 1971; Mosgöller et al., 1991), suggesting that our results could be an indirect effect of different chromosome missegregation positions. To address this hypothesis, we first assessed the location of missegregating chromosomes with different MN stabilities (HSA -1, -11, -17, and -18) during post-mitotic NE assembly, defined as the time between the first appearance of lamin A on anaphase chromatin and the loss of a broad midspindle region, visualized by labeling with alpha-tubulin (Fig. S4A). Our analysis found no significant difference in chromosome missegregation position regardless of chromosome length, gene density, or membrane stability (Fig. 3A – C, Fig. S4B). Second, we analyzed the stability of single chromosome HSA 1 MN when chromosome missegregation was biased towards the spindle pole, by incubation in a CENPE inhibitor (CENPEi), or the midspindle, by incubation in nocodazole (Fig. S4C, D). After a limited arrest (< 6 hours), nocodazole cells were released into fresh medium, and CENPEi cells were released into Mps1i to inhibit mitotic error correction. Our analysis was limited by the fact that only HSA 1 in our pool showed enrichment at spindle poles in CENPEi and that there was only a partial bias towards segregation outside the midspindle after release (Fig. S4C, D). However, no substantial increase was observed for HSA 1 MN stability when midspindle missegregation was reduced (Fig. S4E).

**Figure 3.**
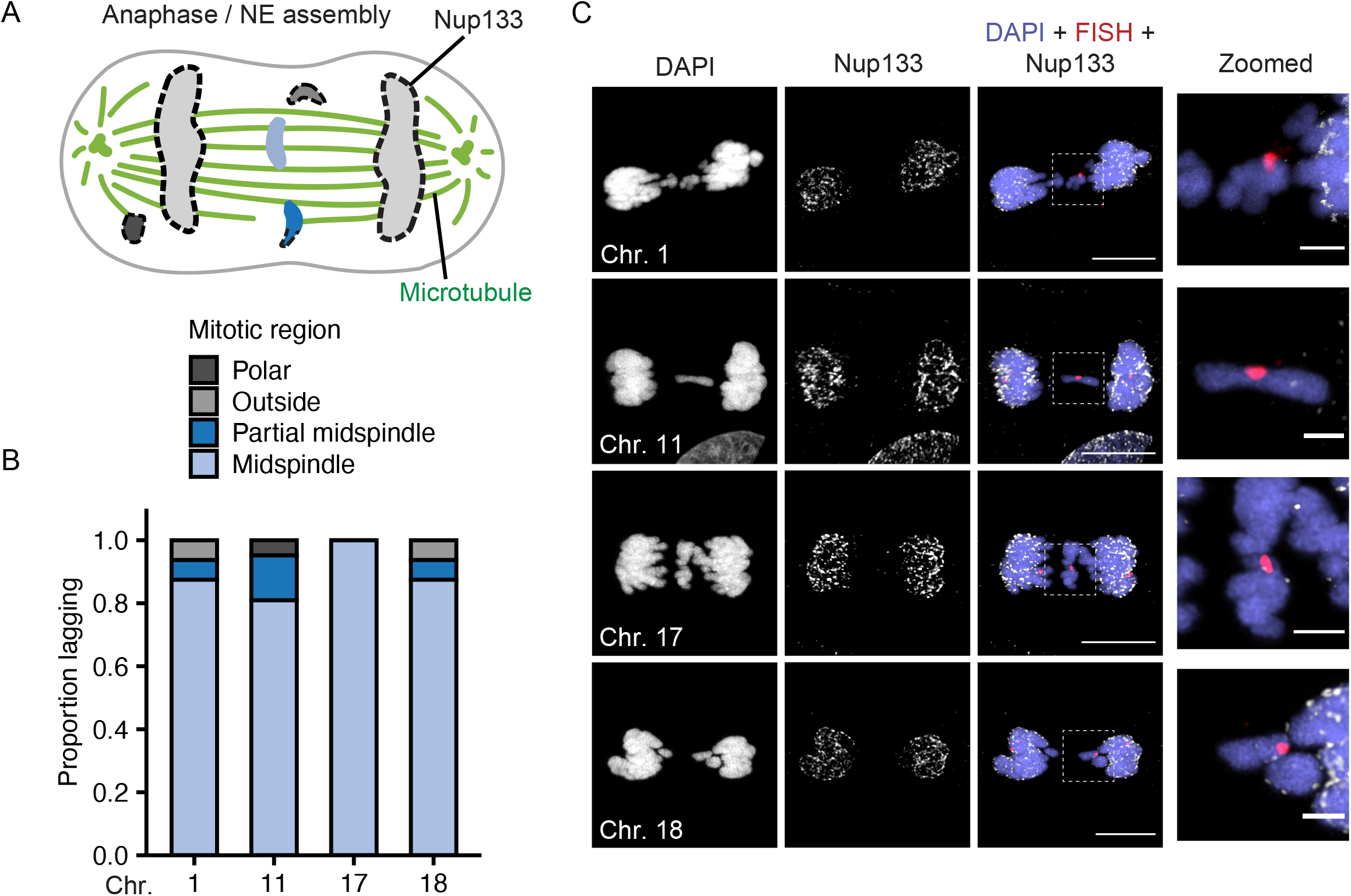
Post-mitotic missegregation position and non-core protein recruitment are not correlated with chromosome identity. (**A**) Model of chromosome missegregation positions and non-core protein (i.e. Nup133) recruitment based on (Liu et al., 2018). Missegregated chromosomes can be within the midspindle (light blue), partially within the midspindle (blue), outside the midspindle (gray), or polar (dark gray). (**B**) Position of indicated missegregated chromosome during early to mid-NE assembly (see Mat. And Meth. for definitions) in RPE-1 cells 13 – 15 hours post-release. N = 3, n = (16, 21, 11, 16). (**C**) Images of Nup133 recruitment to indicated chromosomes during midspindle missegregation. Images are deconvolved single sections (z-step size = 0.15 μm). Scale bar = 10 μm. Zoomed in panels show lower Nup133 foci on all chromosomes compared to main mass. Images are single sections after deconvolution. Scale bar = 3 μm.

Chromosomes missegregating in the midspindle have reduced recruitment of “non-core” proteins, nucleoporins (Nups), lamin B1, and lamin B receptor (LBR), during NE assembly (Afonso et al., 2014; Castro et al., 2017; Karg et al., 2015; Liu et al., 2018). If differences in midspindle association are present between chromosomes, we would expect to observe differences in non-core protein recruitment. To test this hypothesis, we analyzed recruitment of the NPC protein Nup133, an early nuclear pore assembly protein (Otsuka and Ellenberg, 2018), to lagging HSA -1, -11, -17, and -18 chromosomes. Nup133 recruitment was substantially reduced on lagging chromosomes compared to the main chromatin mass, consistent with previous results (Afonso et al., 2019; Liu et al., 2018), and no substantial difference was observed for different chromosomes (Fig. 3C). Together with our observation that chromosome position is not biased during missegregation, these data strongly suggest that chromosome-specific effects on membrane stability are independent of early events in NE assembly.

To assess whether chromosome length and gene density determined MN non-core NE protein levels in interphase, we analyzed lamin B1 levels and NPC density on MN in early G1 (Fig. S1A), to focus on initial protein recruitment and to enrich for intact MN. Lamin B1 intensity was determined using confocal microscopy, and Nup133 foci were quantified using stimulated emission depletion microscopy (STED). As expected, both lamin B1 protein levels and Nup133 foci density were reduced on MN compared to nuclei, though all MN contained at least one Nup133 focus (Fig. 4A – F, S5A, B) (Hatch et al., 2013; Kneissig et al., 2019; Liu et al., 2018; Xia et al., 2019). Surprisingly, lamin B1 intensity and Nup133 density did not correlate with MN stability, but instead with MN size (Fig. 4C,F). Small HSA -17 and -19 MN are significantly more stable than HSA -18 MN (Fig. 1D), yet all three had similar reductions in lamin B1 intensity and Nup133 foci density (Fig. 4B,E). To determine whether the correlation between size and protein levels was limited to single chromosome MN, we analyzed lamin B1 and Nup133 levels in larger MN containing multiple chromosomes. We found that lamin B1 and Nup133 recruitment increased with increasing MN size in multi-chromosome MN as well as single chromosome MN (Fig. 4C,F, S5A, B). Furthermore, we observed that individual chromosomes had a large variance in lamin B1 and Nup133 amounts (Fig. 4B, E), consistent with the variance in MN area observed for specific chromosomes (Fig. 2A). These data strongly suggest that MN size is the main determinant of non-core protein recruitment or maintenance in interphase and that neither lamin B1 nor NPC amount is sufficient to predict membrane stability.

**Figure 4.**
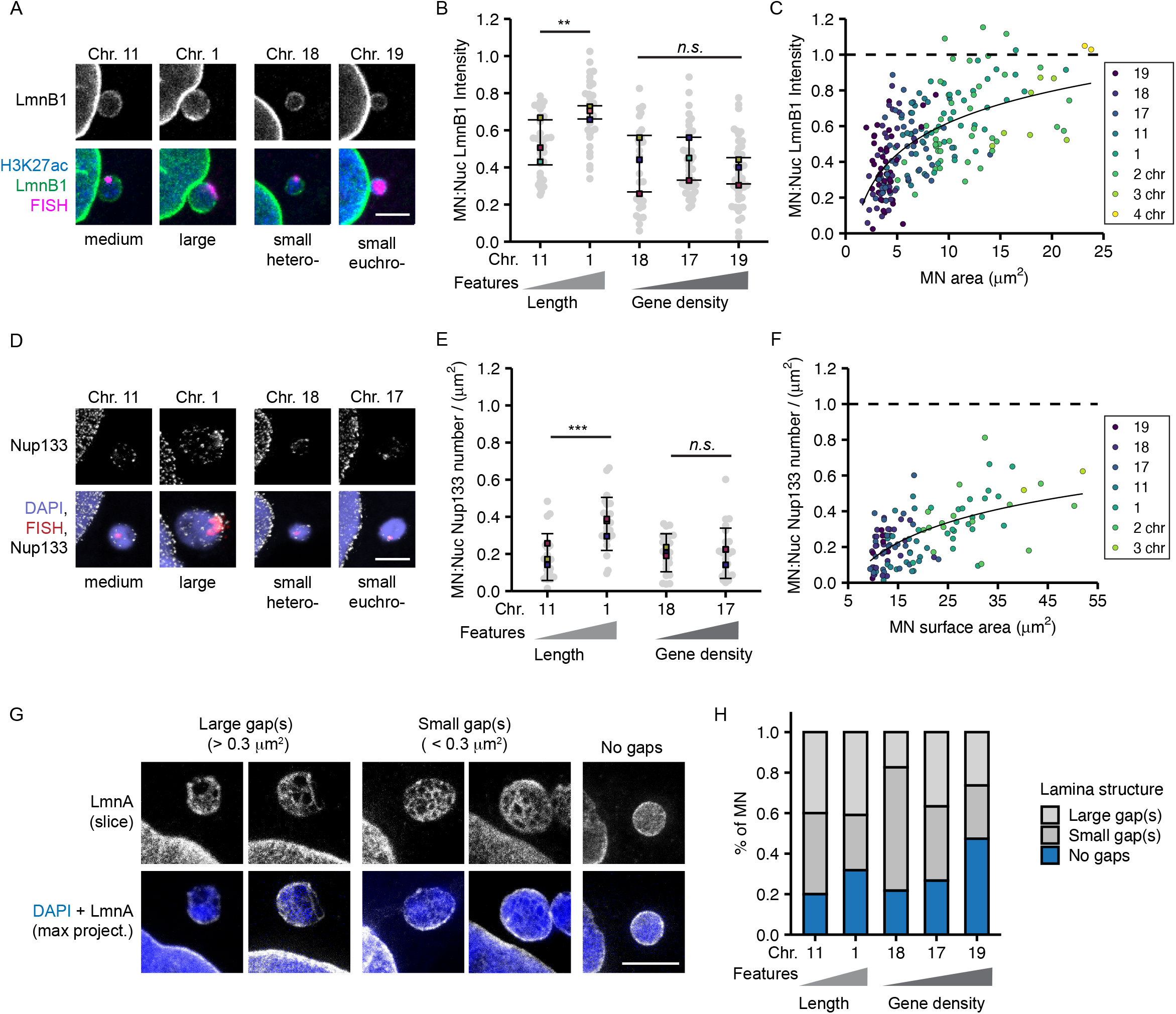
MN size correlates with nuclear lamina protein levels and gene density inversely correlates with nuclear lamina gaps. (**A**) Images of lamin B1 (LmnB1) levels on intact (H3K27ac+) single chromosome MN containing indicated chromosome (see Fig S5A for merged images) 20 h post cdk4/6 release into Mps1i. LmnB1 images are a single section and merged images are maximum intensity projections (MIP). LmnB1 (green), H3K27ac (blue), and FISH (magenta). Scale bar = 3 μm. (**B**) Quantification of MN LmnB1 intensity normalized to nucleus intensity. LmnB1 intensity is increased for longer chromosome MN. ANOVA one-way, p-value < 0.0001; pairwise comparison with Bonferroni adjustment, p-value < 0.01, *n*.*s*. p > 0.05; N = 3, n = (33, 29, 29, 37, 43). (**C**) LmnB1 intensity correlates with MN area for single and multiple chromosome MN Spearman’s correlation (solid line), R = 0.63, p-value < 0.0001. Chromosome number determined by CREST signal (see Fig S5A). Dotted line indicates equal MN and nucleus LmnB1 intensity. N and n for single chromosomes as in B. N = 3 for multi-chromsome MN, 2 chr. n = 41, 3 chr. n = 9, 4 chr. n = 2. (**D**) MIP Images of Nup133 foci on single chromosome MN containing one chromosome rendered in Imaris. MN integrity determined by H3K27ac+ signal (not shown). Scale bar = 3 μm. (**E**) Quantification of Nup133 density (foci/area) on 3D surface normalized to nucleus density LmnB1 intensity is increased for longer chromosome MN. One-way ANOVA, p-value < 0.0001; Bonferroni adjusted pair-wise comparison, *** p-value < 0.001, *n*.*s*. p > 0.05. N = 3, n = (21, 23, 24, 24). (**F**) Nup133 density correlated with MN surface area for single and multiple chromosome MN. Spearman’s correlation (solid line), R = 0.59, p-value < 0.0001. N = 3 for multi chromosome MN, 2 chr. n = 16, 3 chr. n = 2. N and n for single chromosomes as in E, with N = 1 for HSA 19, n = 6 included. (**H**) Representative images of nuclear lamina organization in intact single chromosome MN labeled with antibodies to lamin A (LmnA). MN were classified as containing small nuclear lamina gap(s) when the largest gap was < 0.3 μM ^2^, or large nuclear lamina gap(s) when the largest gap was > 0.3 µm^2^. Top panel images are single sections of MN surface and bottom images are maximum intensity projections. MN integrity determined by H3K27ac labeling (not shown). Scale bar = 3 μm. (**G**) Quantification of nuclear lamina organization in single chromosome intact MN of indicated identity, N = 3, n = (25, 23, 23, 39, 19).

Gaps in the nuclear lamina are thought to be required for nuclear membrane rupture in both nuclei and MN. Lamina gaps are the main sites of membrane blebbing and breakage and occur in response to a variety of conditions, including lamin B1 depletion (Hatch et al., 2013; Hatch and Hetzer, 2016; Maciejowski et al., 2015; Maciejowski and Hatch, 2020; Shimi et al., 2008). To assess nuclear lamina morphology in single-chromosome MN, we imaged RPE-1 cells fixed in early G1 and labeled with antibodies to lamin A using STED. Unlike lamin B1, lamin A is recruited to MN and lagging chromosomes at similar levels to the nucleus, enabling consistent visualization of the lamina meshwork (Castro et al., 2017; Liu et al., 2018). Nuclear lamina gaps were defined as areas with a defined edge that lacked lamin A signal. MN lamina meshwork morphologies were binned into three categories: no gaps, small gap(s) (largest < 0.3 μm^2^), or large gap(s) (largest > 0.3 μm^2^) (Fig. 4G). Examples of MN with small or large gaps were found for all chromosomes and for multi-chromosome MN (Fig. 4H, Fig. S5C). However, qualitative analysis of meshwork morphologies for individual chromosomes identified a consistent trend of increased frequency of MN with no lamina gaps in more stable MN populations, and highly stable MN containing HSA 19 had the fewest lamina gaps (Fig. 4H). These results suggest that large MN size and higher protein density are insufficient to maintain an intact nuclear lamina, and instead an unknown property correlated with high gene density is critical for meshwork organization and membrane stability.

## Discussion

In this study, we demonstrate that chromosome properties are a critical determinant of MN membrane stability. We identify conserved correlations between membrane stability and increased chromosome length and gene density. MN containing a large chromosome or a small, gene dense chromosome rupture later during interphase compared to small gene-poor chromosomes. These correlations cannot be solely explained by differences in chromosome missegregation position or in recruitment of non-core proteins during early NE assembly. Instead, we find that chromosome length, or chromosome number, is directly proportional to MN size and leads to increased levels of lamin B1 and NPC density. High gene density, on the other hand, leads to decreased nuclear lamina gaps, even on small MN depleted of lamin B1 and NPCs. These two mechanisms are independent of each other, as shown by experiments demonstrating that these two features have additive effects on MN stability. Overall, our data support the existing paradigm that nuclear membrane rupture requires nuclear lamina gaps, but strongly suggest that lamin B1 and NPC depletion are insufficient to explain why MN have more nuclear lamina defects compared to nuclei. Instead, we propose that an additional factor, regulated by gene density, determines the appearance of nuclear lamina gaps and the timing of membrane rupture (Fig. 5).

**Figure 5.**
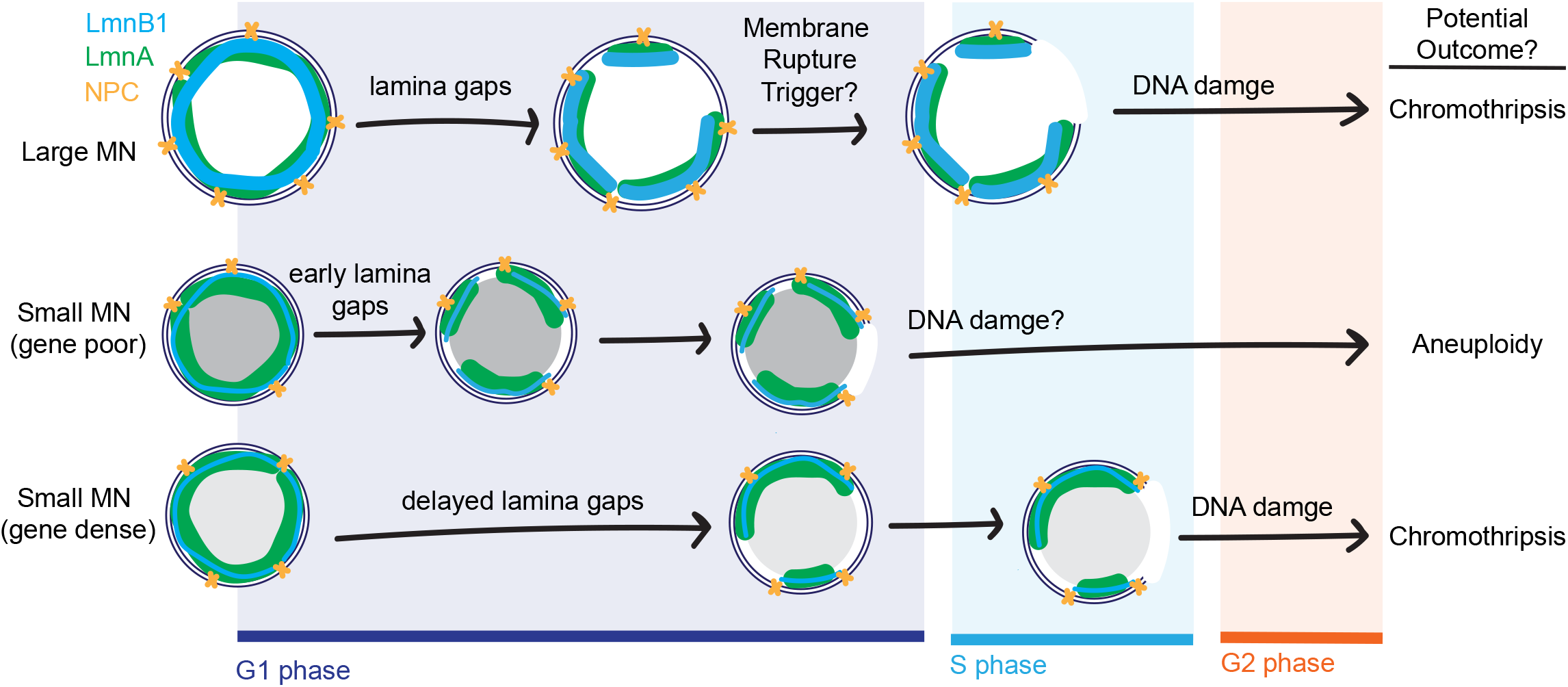
Size and gene density dependent effects on micronuclei lamina structure, rupture timing, and cellular outcomes. The NE is comprised of an inner and outer membrane, NPCs (yellow), and a nuclear lamina made up of A and B type lamins (e.g. LmnA (green) and LmnB1 (blue)). LmnB1 and NPC density depends on MN size regardless of gene density. Despite having high levels of LmnB1, large MN often have gaps in their nuclear lamina. Although gene-dense MN have low levels of LmnB1, they are less likely to contain nuclear lamina gaps compared to gene-poor MN during G1 phase, though gaps are still required for rupture and likely form during late G1. The majority of large and small gene-dense MN rupture during S-phase or later when the DNA is replicating, causing DNA double-strand breaks (DSBs) and chromothripsis. Whether actin forces trigger membrane rupture trigger for single chromosome containing MN is still unclear. The majority of small gene-poor MN rupture in G1 prior to DNA replication, and therefore are less-likely to experience a high level of DSBs but may still experience DNA degradation resulting in aneuploidy.

We observe strong correlations between chromosome length, gene density, and MN stability for multiple mitotic disruptions and in multiple cell lines. However, the average proportion of intact MN differs in different conditions. Although much of this difference could be attributable to differences in cell synchronization efficiency and timing, these observations, along with the high variance in NE protein recruitment to individual micronucleated chromosomes, suggest that chromosome identity is likely one of several factors that determine NE structure and membrane stability. For instance, a recent paper analyzing Y chromosome containing MN in DLD-1 cells found that these MN rarely ruptured during interphase, in contrast to what our results predict for a short, gene-poor chromosome (Ly et al., 2019). Understanding how different conditions affect rupture frequency of specific chromosome MN will be critical to defining these additional factors and how they interact to regulate MN stability.

In the nucleus, nuclear lamina gaps are frequently located at areas of high membrane curvature, and inducing curvature is sufficient to break lamin B1 protein interactions (Deviri et al., 2019; Thiam et al., 2016; Xia et al., 2019). These findings have led to the hypothesis that high membrane curvature in small MN causes loss of lamin B1 leading to lamina gaps (Xia et al., 2019, 2018). However, our observations that MN in cultured cells are oblate spheroids with similar highly curved regions across all size ranges, and that a subset of the smallest MN have a reduced frequency of lamina gaps, argue against a simple curvature model. Although the localization of the largest nuclear lamina gaps to the highly curved edges of the MN (Fig 4G, (Hatch et al., 2013)) suggests that membrane bending promotes rupture. Instead, our results suggest that defects in lamin B1 protein levels and organization arise from a separate mechanism and that this mechanism is active during or after late telophase. At this time, we cannot distinguish between defects in lamin B1 recruitment during NE assembly, import during interphase, or maintenance at the nuclear periphery, which will require chromosome-specific live-cell imaging to unravel.

While gene density is an important factor, it is unclear what aspect of gene density determines nuclear lamina organization. Gene density correlates strongly with GC content, euchromatin histone modifications, transcription, early replication timing, low NE contact, and higher chromatin mobility in interphase (Schneider and Grosschedl, 2007). It is unknown whether functional differences between gene-poor or gene-dense chromosomes are maintained in MN, but it is likely that gene-dense chromosomes have substantially more contact with the NE in MN compared to nuclei (Cremer and Cremer, 2001, 2010; Shimi et al., 2008). Based on our observation that high gene density does not prevent membrane rupture or lamina gap formation (Fig. 1E, Fig. 4H), we hypothesize that one of these characteristics associated with high gene density delays lamina gap formation.

Chromosome size and gene density determine whether the MN is likely to rupture in G1, prior to DNA replication, or after DNA replication initiates (Fig. 1D), and this could have significant effects on DNA damage and the extent of genome instability. Generation of double-stranded DNA breaks in MN both pre- and post-rupture depends on replication initiation (Crasta et al., 2012; Hatch et al., 2013; Umbreit et al., 2020; Zhang et al., 2015), whereas the cytoplasmic nuclease TREX1 (three prime repair exonuclease 1) likely acts on ruptured MN throughout the cell cycle to generate ssDNA (Mohr et al., 2021). Currently, only MN that remain intact until NE disassembly or rupture *after* initiating DNA replication have been shown to undergo chromothripsis (Ly et al., 2019, 2016; Umbreit et al., 2020; Zhang et al., 2015). In contrast, early MN rupture causes whole chromosome aneuploidy in interphase and the next cell cycle. In addition, TREX1 inhibits cGAS-STING activation, suggesting that rupture timing could affect innate immune signaling as well (Harding et al., 2017; Mackenzie et al., 2017; Mohr et al., 2021). How these two pathways of DNA damage, and chromosome-specific differences in transcription, replication timing, and NE assembly interact to determine the consequences of micronucleation and MN rupture remains to be determined. Identifying the chromosomes that missegregate into MN in different tissues and cancer types will be critical to understanding how MN rupture drives cancer evolution and disease pathogenesis *in vivo*.

## Materials and Methods

### Cell lines and culture methods

hTERT-RPE-1 (RRID: CVCL_4388) cells were grown in DMEM/F12 (Gibco) + 10% FBS (Gibco) + 1% Pen-Strep (Gibco) + 0.01 mg/ml hygromycin (Sigma-Aldrich) at 5% CO2 and at 37°C. hTERT-RPE-1 3xGFP-NLS is a stable cell line characterized previously (Anderson et al., 2009). IMR90 cells (RRID: CVCL_0347) were grown in DMEM (Gibco) + 15% FBS (Gibco) +1% Pen-Strep (Gibco) at 5% CO2, 5% O2 at 37°C. Cell line identity was determined by short tandem repeat typing.

For RPE-1 FISH experiments, except where noted, cells were arrested in G1 by addition of 1 uM PD-0332991 isethionate (Sigma-Aldrich) for 24 hours. Cells were released by washing three times in 1x PBS prior to incubation in medium containing the Mps1 inhibitor reversine, (0.5 μM or 1 μM; EMD Millipore) or BAY-1217389, (100 nM; Fisher) for 14 to 32 hours. For nocodazole treatment and missegregation position experiments, RPE-1 cells were incubated in 100 ng/mL nocodazole (Sigma-Aldrich) or 50 nM GSK-923295 (Cayman Chemicals) for 4 to 6 hours prior to release by washing three times with 1x PBS and then adding either media alone (nocodazole) or media + 0.5 μM reversine (GSK-923295). For mitotic shake off experiments, cells were shaken off after last 1x PBS wash and fixed either during anaphase or 8 hours post mitosis. IMR90 cells were incubated for 32 hours in 1 μM PD-0332991 isethionate then released into 0.5 μM reversine for 24 hours.

### Immunofluorescence (IF) and Fluorescence *in situ* hybridization (FISH)

Cells were grown on poly-L-lysine coated coverslips and fixed in 4% paraformaldehyde (PFA) (Electron Microscopy Sciences) for 10 min at room temperature (RT) for the following experiments: Mps1i total MN rupture frequency time course, MN area to volume analysis, experiments using hTERT-RPE-1 3xGFP-NLS, and multi-chromosome MN analysis using a PNA CENPB-Cy5 probe. For all other IF and FISH experiments, cells were fixed in 100% methanol at -20°C for 5 – 10 min. Coverslips were blocked in 3% BSA (Sigma) + 0.1% - 0.4% Triton X-100 (Sigma-Aldrich) + 0.02% sodium azide (Sigma-Aldrich) in 1x PBS (PBS-BT) for 30 min prior to incubation in primary antibodies diluted in PBS-BT. Primary antibodies used: mouse alpha-tubulin (1:250; 3873S, Cell Signaling Technology), human anti-CREST (1:100; 15-234, Antibodies Incorporated), mouse anti-CENPA (1:100; GTX13939, GeneTex), mouse or rabbit anti-H3K27ac (1:250; 39085, Active Motif; 1:1000; ab4729, Abcam), rabbit anti-Lamin B1 (1:100; sc-365214, Santa Cruz Biotechnology), rabbit anti-Lamin A (1:500; L1293, Sigma-Aldrich), rabbit anti-Nup133 (1:100; ab155990, Abcam). Coverslips were washed three times in PBS-BT then incubated in the following secondary antibodies: Alexa Fluor 405-conjugated goat anti-rabbit (1:2000; A-31556, Thermo Fisher Scientific), Alexa Fluor 488–conjugated goat anti–mouse (1:2000; A-11029, Thermo Fisher Scientific), Alexa Fluor 488–conjugated goat anti–rabbit (1:2000; A-11034, Thermo Fisher Scientific), Alexa Fluor 594–conjugated donkey anti-rabbit (1:500; 711-585-152; Jackson ImmunoResearch), Alexa Fluor 647–conjugated goat anti–mouse (1:1000; A-21236; Thermo Fisher Scientific), Alexa Fluor 647–conjugated goat anti-human (1:1000; A-21445; Thermo Fisher Scientific). Secondary antibodies were dilute in PBST and incubated for 30 minutes at RT. Coverslips were washed twice in PBST then incubated with DAPI (1 μg/ml in PBS; Roche) for 5 min at RT, washed once in diH_2_O, and mounted in Vectashield (Vector Labs) or Prolong Gold (Life Technologies).

For experiments using chromosome enumeration (XCE) or whole chromosome paint (XCP) probes. after methanol fixation and immunofluorescence as described above up until DAPI labeling, coverslips were fixed for 5 minutes with 4% PFA in 1x PBS. This and subsequent steps were performed at RT unless noted. Coverslips were washed twice with 2x SSC (Sigma-Aldrich) for 5 min then permeabilized with 0.2 M HCl + 0.7% Triton X-100 for 10 – 15 min at RT. Coverslips were washed twice with 2x SSC for 5 min, denatured in 50% formamide (EMD Millipore) 2x SSC for 1 hour, washed twice with 2x SSC, then inverted onto 3 – 5 μL of Spectrum Orange XCE or XCP probe (MetaSystems) and sealed with rubber cement. Probes and targets were co-denatured at 74°C for 2 min and hybridized 2 hours to overnight at 37 °C in a humidified chamber. Coverslips were washed once in pre-heated 0.4x SSC buffer at 74°C for 5 min then twice in 4x SSC or 2x SSC + 0.1% Tween-20 for 5 min. Coverslips were incubated in DAPI and mounted in Vectashield (Vector Labs) (for analysis of MN morphology or protein recruitment) or Prolong Gold (Life Technologies).

For FISH with the PNA CENPB-Cy5 probe, the same protocol was followed until formamide denaturation. At that point cells were incubated in 50% formamide in 2x SSC for 30 min at 85°C then rinsed three times in ice cold 2x SSC. PNA probes were diluted to 50 μM in 85°C hybridization buffer (60% formamide + 20 mM Tris, pH 7.4 + 0.1 μg/mL salmon sperm DNA (Trevigen)) and coverslips were simultaneously washed at 85°C in 2x SSC. Coverslips were then incubated in 10 μL of the PNA probe for 10 min at 85°C and then 2 hours at RT. Coverslips were then washed twice with 2x SSC + 0.1% Tween-20 at 55°C for 10 min and once with 2x SSC + 0.1% Tween-20 at RT. before incubation in DAPI and mounting in Prolong Gold (Life Technologies).

### EdU-pulse labeling and fluorescence activated cell sorting (FACS)

For FACS cell cycle analysis, cells were incubated with 10 μM EdU (Life Technologies) for 15 min in culture media prior to trypsinization and fixation in 70% ice cold ethanol. Cells were stored at -20°C prior to staining. Fixed cells were washed twice in 1xPBS + 0.1% Triton-X 100 before resuspension in Click-It EdU reaction mix (ThermoFisher; Alexa-555) for 30 minutes while rotating. Cells were washed twice in 1xPBS + 1% BSA and incubated in 1 ug/mL DAPI in 1x PBS for 30 minutes prior to analysis. Samples were analyzed on either a 3-laser FACSCanto™ II (BD Biosciences) or a 4-laser LSR II (BD Biosciences) and data acquired using DIVA software (BD Biosciences). DNA content was analyzed based on DAPI fluorescence, and DNA replication was analyzed based on Alexa-555 fluorescence. Doublet discrimination was used to remove doublets and clumped using DAPI-A/DAPI-W measurements. DAPI and Alexa-555 signal were plotted as area measurements. Data were analyzed using FlowJo v.10 software (BD Biosciences). Cell cycle distributions were determined by gating EdU positive versus negative, as determined by single color control, and by 2N versus 4N DAPI content.

### Microscopy

Confocal images were acquired with a Leica DMi8 laser scanning confocal microscope using the Leica Application Suite (LAS X) software and with the Leica ACS APO 40x/1.15 Oil CS objective or a Leica ACS APO 63x/1.3 Oil CS objective. Z-stacks were acquired with the system optimized step size or at 0.5 μm for IMR90 and mitotic images. Confocal images of mitotic cells in figures 3C and D were acquired with a Leica TCS SP8 confocal microscope with a pixel size between 60 – 80 nm and a z-step size of 0.15 um with a Leica HCX Plan Apo 63x/1.40 Oil CS2 objective. Post-acquisition, images were deconvolved using Lightning with smoothing size of medium through the LAS X software.

Nup133 and lamin A IF labeled cells were imaged using Leica TCS SP8 with the super-resolution microscope system (STED) using a 775 nm pulsed laser, Leica Application Suite software platform (LAS X version 3.5.7.23225), and a Leica HC PL APO 100x/1.4 Oil CS2 objective. Prior to image acquisition the STED and confocal beams were manually aligned using FluoSpheres mounted in Prolong Gold and white light laser set to 594 nm and 775 nm STED, the alignment was adjusted until the STED FluoSpheres overlaped with the center of the confocal FluoSpheres images. Images were acquired at ∼20 nm pixel size for a resolution of approximately 50 nm in the xy plane, and a white light laser was tuned to 405 nm (DAPI), 488 nm (H3K27ac), 556 nm (FISH), 594 nm (Nup133 or lamin A), 647 nm (CREST) wavelengths.

Post-acquisition image processing was limited to cropping the image and adjusting levels through Adobe Photoshop to make use of the entire histogram spectrum. False colors for channels were changed through the arrange channels function in Fiji (Schindelin et al., 2012).

### Image Quantification

MN were defined as a DAPI positive round object adjacent to or near the nucleus that was distinct from the nucleus to distinguish them from nuclear herniations, and chromatin bridge fragments. Teardrop shaped objects were excluded from analysis. Intact MN were defined as those with H3K27ac signal that was of the same average intensity as the H3K27ac signal of the main nucleus over some part of its area. Ruptured MN were defined as those where the average H3K27ac signal was decreased by > 60% compared to the main nucleus. Chromosome number was defined as the number of centromere foci, which were assessed by CENPA or CREST IF, or PNA CENPB-Cy5 FISH. A positive FISH signal was defined as a focus twice the signal of background that partially co-localized with a centromere. Interphase cells with more than three FISH foci were excluded from analysis as being either tetraploid or exceeding acceptable signal to noise ratios.

MN area was calculated from maximum intensity projections by selecting the DAPI channel object and measuring the area in Adobe Photoshop. Pixel area was then converted to μm^2^ based using the image calibration dimensions.

Missegregated chromosomes in mitotic positioning analysis were defined as FISH positive chromsomes that were not contiguous with the main chromatin mass during early-mid nuclear envelope assembly (NEA). This stage was defined by the presence of a wide spindle midzone and recruitment of lamin A to the main chromatin mass in either a punctate or continuous pattern. These conditions were chosen based on previous work demonstrating that nuclear import starts after the initiation of cytokinesis in RPE-1 cells.

LmnB1 intensity was quantified for intact (H3K27ac+) MN containing a single CREST focus that overlapped with the FISH probe 20 hours post-cdk4/6i release into 0.5 μM MPS1i. A single z-slice was analyzed from the middle of the MN and corresponding nucleus and the average intensity of the entire rim was taken for each MN to account for patches of dim signal that likely corresponded to lamina gaps. Images were imported into Adobe Photoshop and the quick selection tool was used to outline the nuclear perimeter of the MN and nucleus from the H3K27ac channel. A four pixels border was added to the perimeter outline to encompass the entire LmnB1 rim signal. Measurements were taken for the perimeter (pixels), border area (pixels), and integrated density. The background was subtracted for both the nucleus and MN using a border expansion of 5 pixels. The background-subtracted mean fluorescence intensity was calculated by dividing the background subtracted integrated density from the 4 px wide boarder area.

Nup133 density was determined for intact (H3K27ac+) MN containing a single CREST focus that overlapped with the FISH probe 20 hours post-cdk4/6i release into 0.5 μM MPS1i. The number of Nup133 foci for each MN and nucleus were quantified in Imaris x64 8.4.2 by first defining the region of interest (ROI) around each MN and nucleus using the contour tool, then creating spots for each ROI with a XY spot diameter set to 0.2 μM. The threshold was adjusted for each image to capture every Nup133 focus in the nucleus but very few spots in the cytoplasm. The same threshold was used for the corresponding MN. The surface area was calculated in Imaris from the DAPI channel, with smoothing set to 0.5, and background subtraction of 0.2, and the threshold adjusted to encompass the entire DAPI signal.

Nuclear lamina gaps were defined as areas with a defined edge that lacked lamin A signal and meshwork morphologies were binned into three categories: none (no gaps > 0.12 μM ^2^), small gap(s) (largest gap between 0.12 – 0.3 μM ^2^), or large gap(s) (largest gap > 0.3 µm^2^). MN lamina gap area for the largest gap were measured in FIJI.

### Statistics

Analysis of two group categorical data were conducted using Barnard’s exact test, run on either MATLAB (version 2018b) Barnardextest (verson 1.0.0.0) or R (version 4.0.0) using the ‘Barnard’ package. All additional statistic tests were conducted using either Prism 8 or R (version 4.0.0). Statistical analysis of two-group quantitative data were performed using Welch unpaired t-tests. A family test (i.e. Chi-squared for categorical data, ANOVA or Kruskal-Wallis one-way test for continuous data) was first performed for experiments with three or more groups, and only significantly different dataset were analyzed by multiple comparison testing. One-way ANOVA test was used for statistical analysis of quantitative data with normal distribution containing two or more groups followed by Bonferroni corrected pair-wise comparison. Kruskal-Wallis one-way analysis was used to compare three of more independent groups when the assumptions of one-way ANOVA (i.e normality and equal variance) were not meet. Significant association between two binary variables (i.e. H3K27ac and GFP-NLS) was analyzed using the phi mean squared contingency coefficient. Spearman’s rank correlation coefficient was used to assess montonic relationship for two variables. Spearman’s was used for MN area to volume correlation due to the non-normal distribution determined by the Shapiro-Wilk test for normality. For all tests, p-values greater than 0.05 were considered statistically significant. A limitation of chi-square test is that it is highly sensitive to sample size, therefore, a post-hoc analysis was performed to determine if our datasets reach a Chi-square statistical power of 0.8 based on a given effect and sample size using the packages ‘esc’ and ‘pwr’ in R (version 4.0.0). Post-hoc power for lamin A gap proportion (Fig. 4I) analysis yielded a statistical power = 0.595 given an effect size d = 0.582 and N = 3, n = (25, 23, 23, 39, 19). Chi square statistical tests cannot be performed on datasets with 0 values and are invalid when multiple outcomes have a value less than 5, and analysis were not performed on experiments where this was the case.

## Acknowledgments

E.M.H, A.L.G., E.C., and H.Z.H. were supported by the National Institutes of Health grant R35GM124766 and a Rita Allen Foundation Scholars Award. This work was supported by the Cellular Imaging and Flow Cytometry Shared Resources of the Fred Hutch/University of Washington Cancer Consortium (P30 CA015704). A.E.M. was supported by the National Cancer Institute of the National Institutes of Health training grant T32CA009657.

## Notes

### Competing Interest Statement

The authors have declared no competing interest.

