## Supplemental figure legends for "Chromosome length and gene density contribute to micronuclear membrane stability"

**Figure S1.** H3K27ac is a reliable marker of micronuclei membrane integrity (**A**) 94.8% of H3K27ac+ micronuclei were 3xGFP-NLS+; 7.8% of H3K27ac+ micronuclei lacked 3xGFP-NLS-. N = 3 replicates; n > 30 MN; 0.87 phi correlation coefficient; **** p-value < 0.0001. (**B**) Image of an RPE-1 cell containing one intact (top; GFP-NLS+, H3K27ac+) and one ruptured (bottom; GFP-NLS -, H3K27ac -) MN. MN outlined in GFP-NLS (green), H3K27ac (red), and merge images. (**C**) Method for inducing MN in RPE-1 cells and fixation timepoints 20, 24, or 30 hours after Cdk4/6i release into Mps1 inhibitor (Mps1i). (**D**) Approximately 50% of MN are ruptured by G1/S phase as determined by H3K27ac signal. N = 3, n > 100 MN per replicate. Chi-square; p-value > 0.0001. (**E**) Example images of intact single chromosome containing MN (circle) and corresponding nuclei in RPE-1 cells 24 hours post-release. HSA -1, -11, -15, -17, and -20 were probed using spectrum orange XCE probes. HSA -4, -13, -18, -19, and -22 were probe with spectrum orange XCP probes. MN containing chromosomes are indicated by dotted circles. (**F**) Fixed chromosome features: Karyotype refers to chromosome size and centromere positioning. Chromosome length in megabases (Mb) corresponded to human genome assembly GRCh38.p13. Gene density from (Worrall et al., 2018). Centromere length in Mb from human centromere assembly GCA_000442335.2 (Miga et al., 2014). Scale bars = 10 μm. (**G**) Dual parameter plots of RPE-1 EdU-incorporated cells 20, 24, and 30 hours post-Cdk4/6i release into 0.5 μM Mps1i. Gating used to determine the percentage the cells in G1, S, and G2 phase are shown (black boxes). (**H**) Mean proportion of RPE-1 cells in G1, S, and G2 phase per treatment condition. N = 3, n = 6000.

**Figure S2.** Relationship between chromosome features and micronuclei membrane stability. (**A**) Centromere size does not have a strong positive or negative correlation with MN rupture frequency. (**B, C**) Centromere position and rDNA repeats do not significantly impact MN membrane stability when controlled for (B) gene density or (C) chromosome size. N = 3 – 4 replicates, n > 65 MN. (B) MN stability of acrocentric / rDNA containing chromosomes correlates with gene density. (**D**) Enhanced MN stability associated with chromosome length and gene density observed with Mps1 inhibitor (BAY-1217389) treatment. HSA -1 and -11 stability did not differ significantly (Chi - square power = 0.73); N = 3 – 5, n = (86, 82, 99, 83, 81). Barnard's exact test used for all comparisons; * p < 0.05, ** p < 0.01, *n.s.* p > 0.05.

**Figure S3.** Chromosome number regulates MN size and rupture timing. (**A**) Very strong correlation between max projected MN area ($\mu m^2$) and volume ($\mu m^3$). Spearman's correlation, R = 0.904, **** p-value < 0.0001; n = 48. (**B**) Example images of a ruptured (H3K27ac -) MN lacking CENPB signal and intact (H3K27ac+) MN containing 1 – 5 CENPB foci. MN perimeter outlined in CENPB channel. Scale bar = 10 μm. (**C**) Significantly decreased MN stability at various points during interphase, 20, 24, 28, and 32 hours post-Cdk4/6i release into 1 μM Mpsi. for MN containing one or two chromosomes (1 CENPB or 2 CENPB foci), n = (181, 176, 198, 162) and n = (50, 50, 51, 46), respectively. MN containing 3+ chromosomes trended towards reduced stability overtime but did not reach significance; p-value > 0.05, n = (29, 56, 22, 15). Chi-square test, * p < 0.05, **** p-value < 0.0001. N = 3 replicates.

**Figure S4.** Missegregated chromosome positioning during mitosis. (**A, B**) RPE-1 released into 0.5 μM Mps1i for 15.5 hours. (**A**) Cells in anaphase / NE assembly were determined by lamin A (LmnA, gray) and α - tubulin (green) staining consistent with active NE assembly. Individual chromosomes were identified using FISH (magenta) shown with DAPI (gray or blue). Images taken using Leica DMi8 confocal. In all images lamin A is present on the chromatin and the midspindle region has not been collapsed by cytokinesis. (**B**) Uncropped, unrotated images from (Fig. 3C) showing chromosome missegregation (FISH, magenta) within the α–tubulin (green) dense region of the midspindle. Single z-sections are shown (i.e. slice), which contain the lagging chromosome. Scale bar = 10 μm. (**C**) Quantification of HSA 1 and total chromosome missegregation position 45 minutes after release from indicated drug treatment. Chromosome position determined by colocalization of DAPI signal with α-tubulin in midspindle. Partial midspindle localization was scored as "midspindle" in this experiment. No significant difference between the proportion of HSA 1 in each region compared to all chromosomes post CENPEi (GSK-923295) shake-off. All chromosomes including HSA 1 missegregated in the interior mitotic spindle following nocodazole shake-off. Barnard's exact test; *n.s* $p > 0.05$, N = 3, n = (258, 30, 118, 32). (**D**) Images of missegregated single HSA 1 chromosomes after 6 hours incubation in either CENPEi (GSK-923295) or nocodazole (Noco.) and release into 0.5 mM Mps1i or fresh medium, respectively. Scale bar = 10 μm. (**E**) MN stability quantified 8 h post mitotic shake-off and release from indicated drug treatments. As in D, CENPEi-treated cells were released into 0.5 mM Mps1i. Barnard's exact test; *n.s.* $p > 0.05$, N = 3, n = (85, 84). All scale bars = 10 μm.

**Figure S5.** Nuclear envelope composition of micronuclei containing one or multiple chromosomes. (**A**) Lamin B1 (LmnB1, green) signal for a single z-section taken from the center of the nucleus and intact (H3K27ac+, blue) MN. Merged MIP images shown below, H3K27ac (blue), FISH (red), and CREST (gray). (**B**) Imaris rendered images of Nup133 (gray) signal for the nucleus and intact MN (H3K27ac not shown). (**C**) Example of an intact (H3K27ac+) MN containing two chromosomes and several lamin A (LmnA, gray) gaps. The top three Z-slices were max-projected for LmnA, and the merged images of MIP for DAPI (blue), H3K27ac (green), CREST (red). FISH signal not shown for multiple chromosomes containing MN. Scale bars = 10 μm, and 3 μm (zoomed images).
