## Supplemental figure 1 for "Chromosome length and gene density contribute to micronuclear membrane stability"

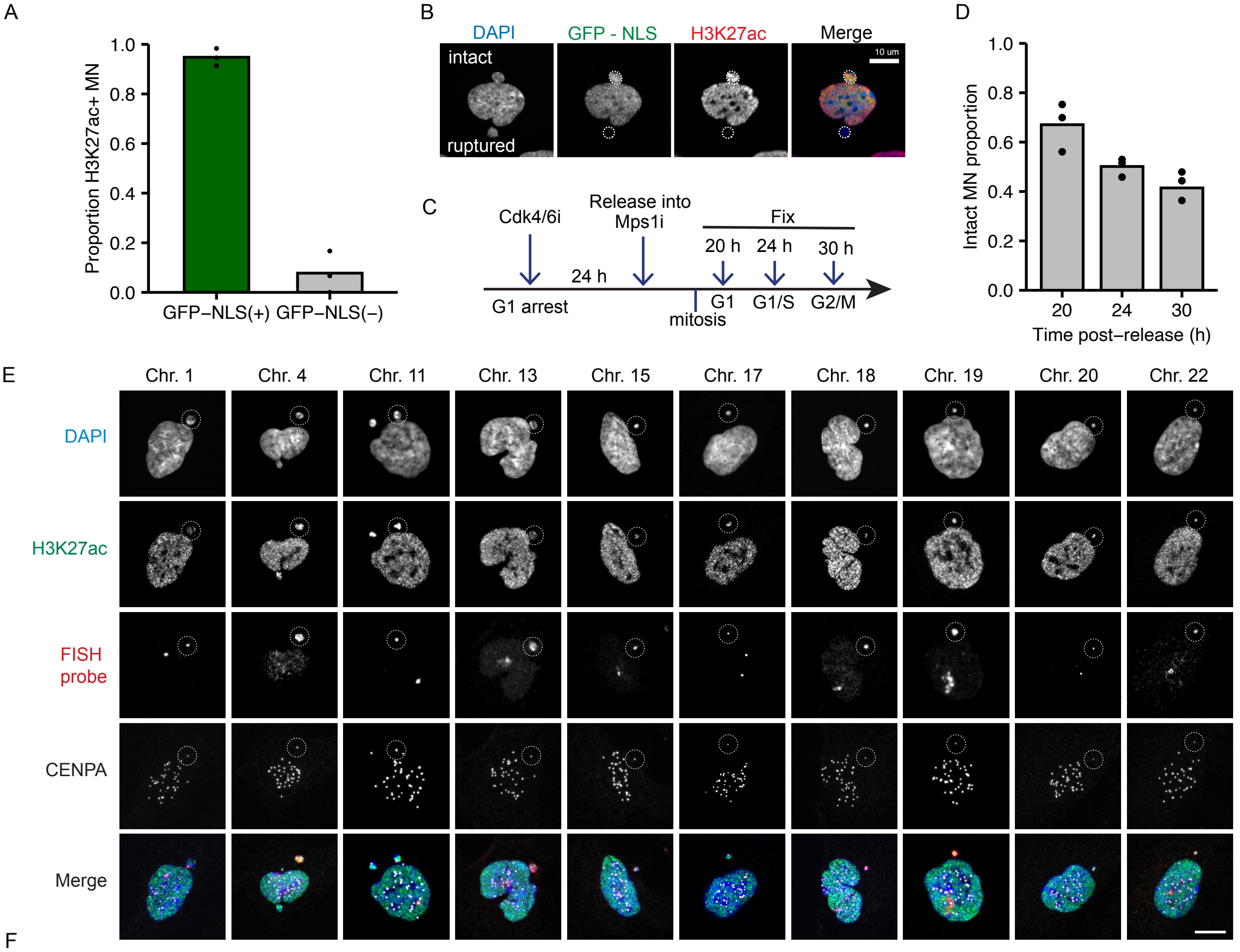


F

| Chromo. features | Chr. 1 | Chr. 4 | Chr. 11 | Chr. 13 | Chr. 15 | Chr. 17 | Chr. 18 | Chr. 19 | Chr. 20 | Chr. 22 |
|---|---|---|---|---|---|---|---|---|---|---|
| Karyotype | Large Metacentric | Large Sub - meta. | Medium Sub - meta. | Medium Acrocentric | Medium Acrocentric | Small Sub - meta. | Small Sub - meta | Small Metacentric | Small Metacentric | Very Small Acrocentric |
| Length (Mb) | 249.0 | 190.2 | 135.1 | 114.4 | 102.0 | 83.3 | 80.4 | 58.6 | 64.4 | 50.8 |
| Gene Density (Genes/Mb) | 20.31 | 12.85 | 21.63 | 12.11 | 17.78 | 29.30 | 12.35 | 42.05 | 21.08 | 22.98 |
| Cent. Length (Mb) | 3.16 | 2.04 | 3.35 | 2.05 | 2.73 | 4.07 | 5.40 | 2.69 | 3.58 | 2.10 |

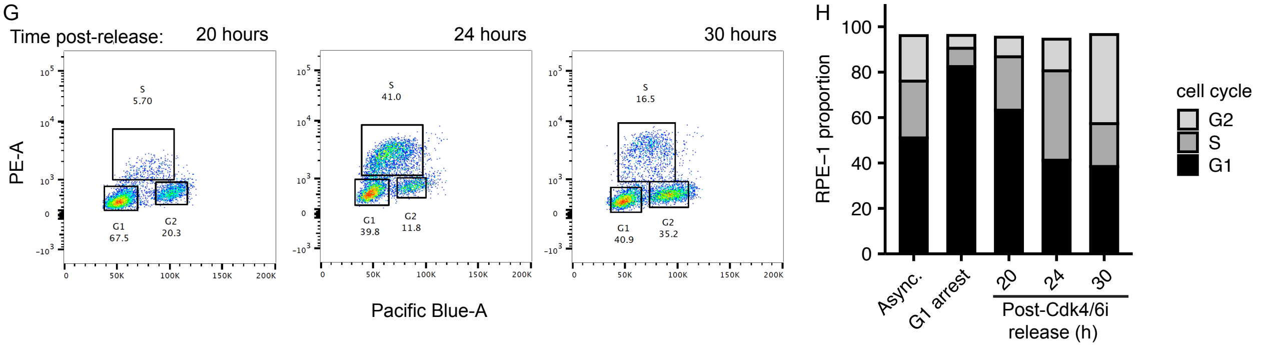
