## Supplementary figures and images for "Chromosome length and gene density contribute to micronuclear membrane stability"

### Supplemental figure 2

#### Supplemental Figure 2

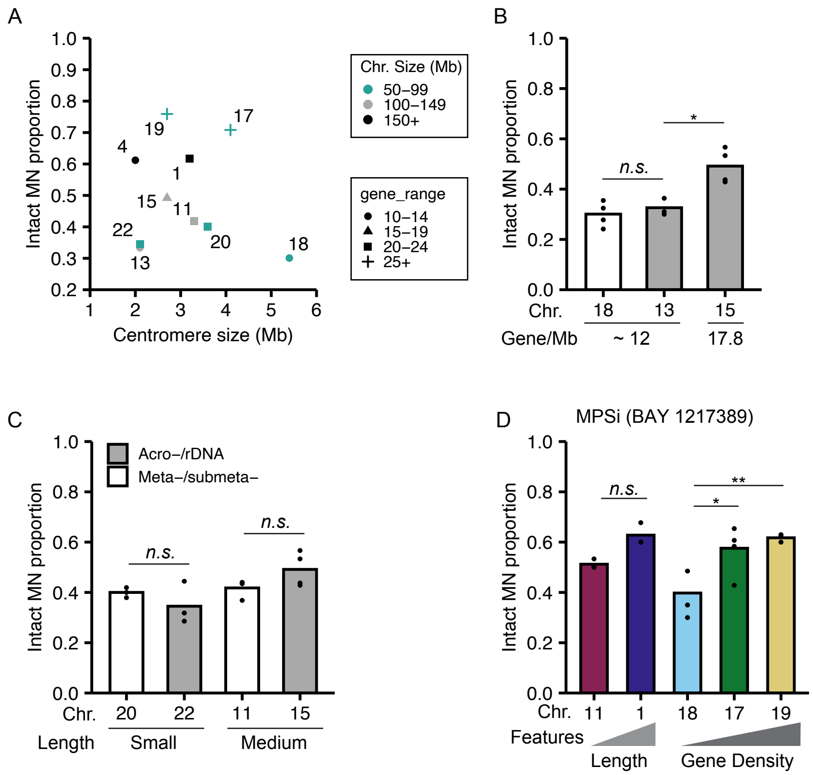

### Supplemental figure 3

#### Supplemental Figure 3

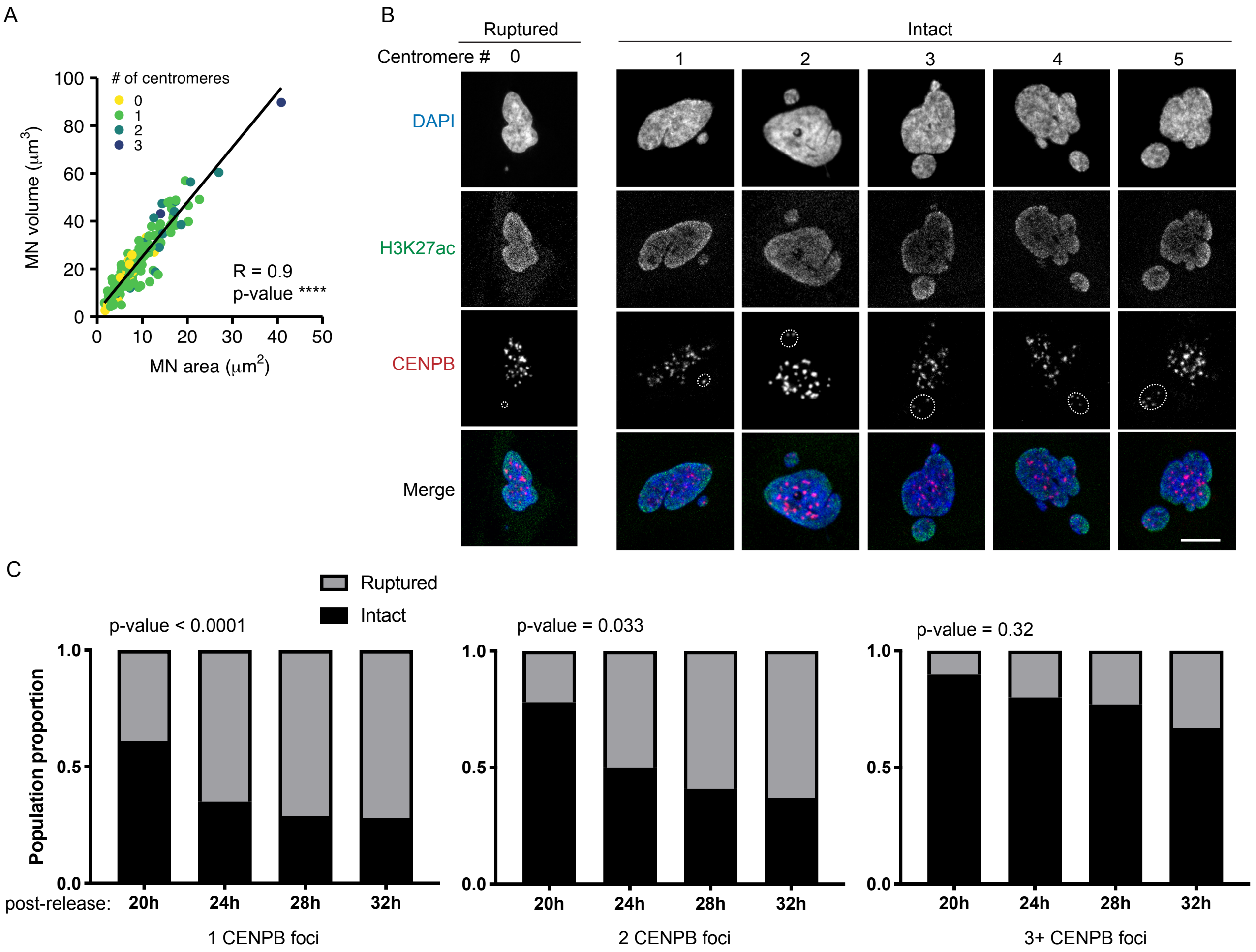

### Supplemental figure 4

#### Supplemental figure 4

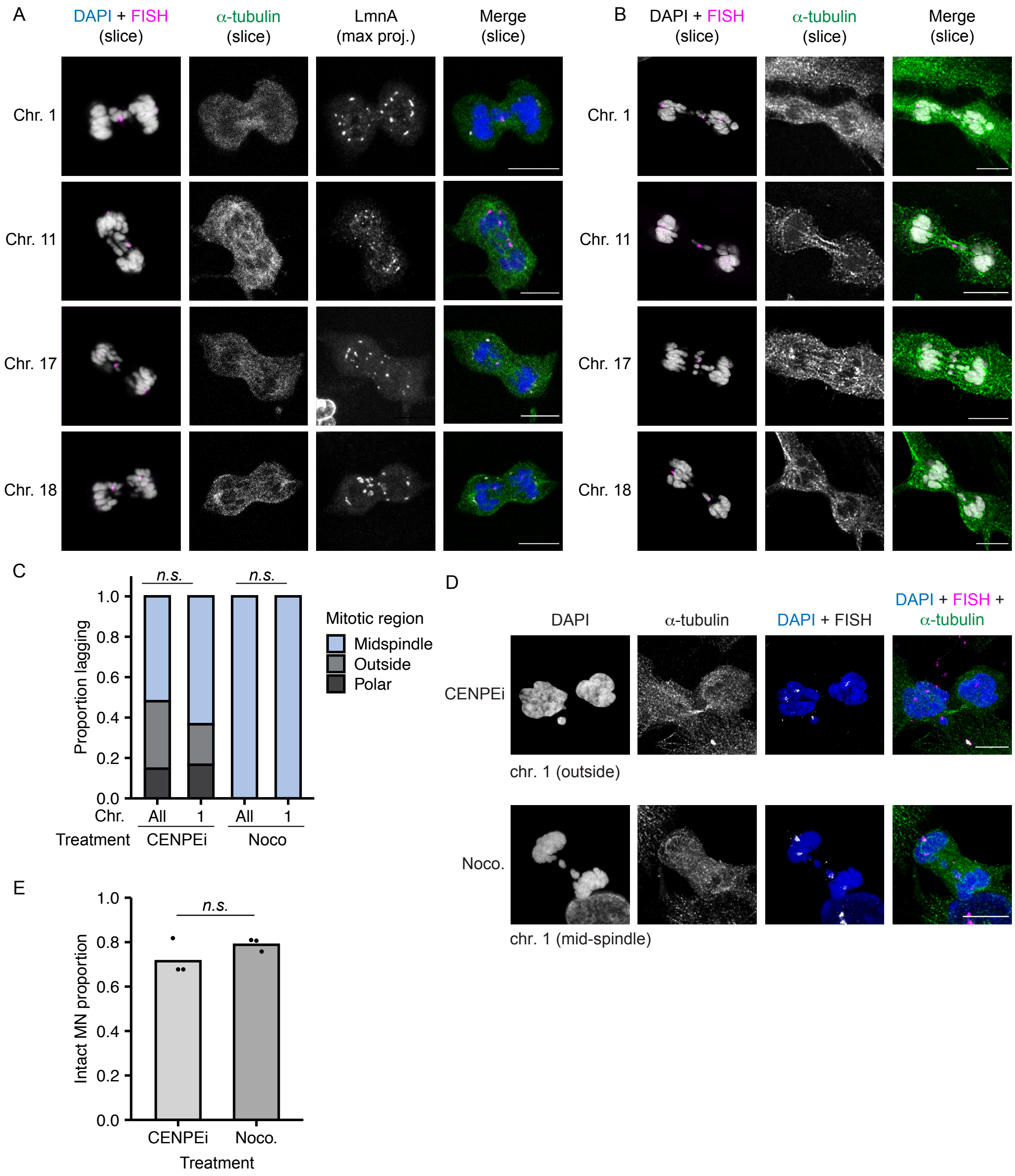

### Supplemental figure 5

#### Supplemental figure 5

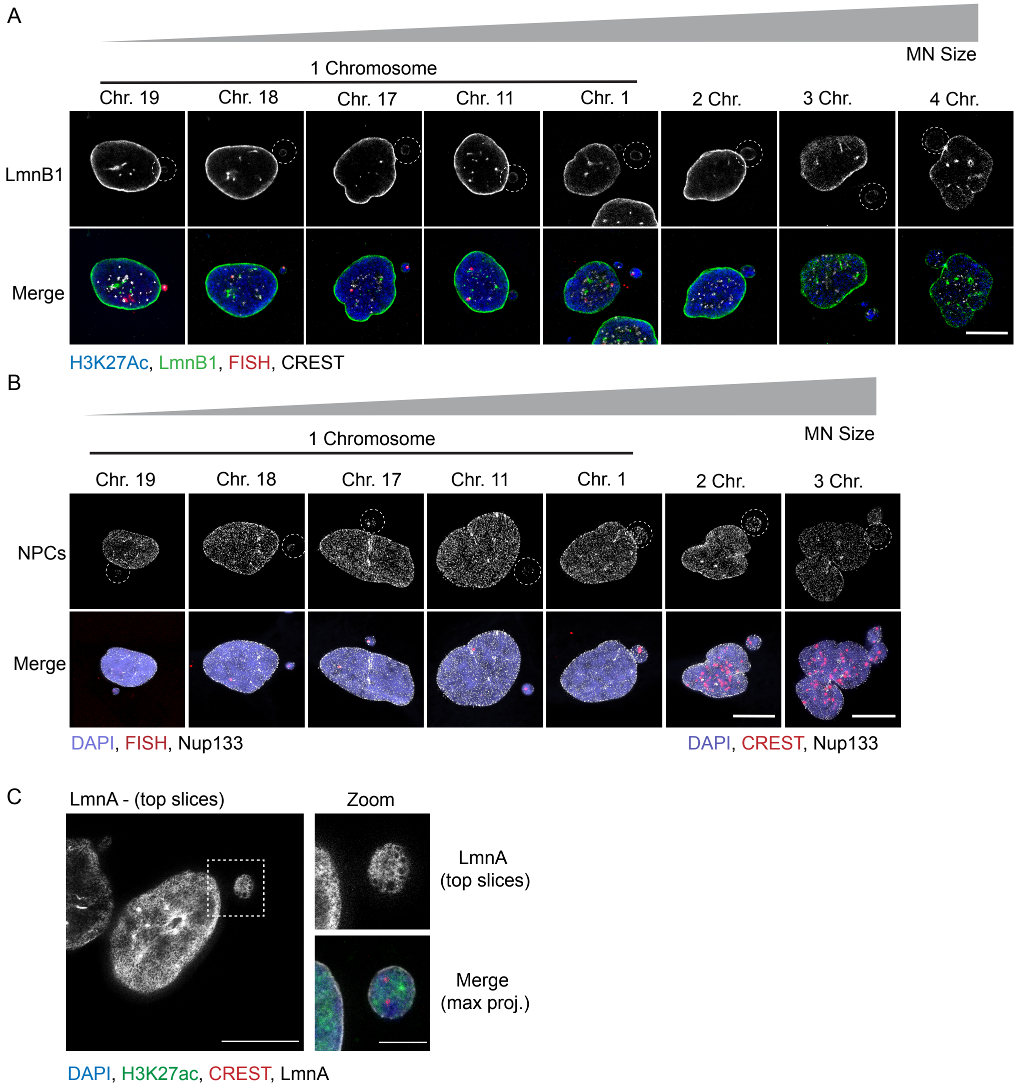
